# Early-life optimization of mitochondrial aerobic metabolism: high efficiency to grow fast, yet at no oxidative cost

**DOI:** 10.64898/2026.04.14.718352

**Authors:** Lucie Lenoel, Damien Roussel, Jessica Barbe, Angeline Clair, Laetitia Averty, Ludovic Calandreau, Antoine Stier

## Abstract

Variation in mitochondrial aerobic metabolism has been suggested to underlie variation in interindividual performance. Mitochondrial efficiency quantifies, directly or indirectly, the amount of adenosine triphosphate (ATP) produced relative to O_2_ consumption. High mitochondrial efficiency is theoretically beneficial by providing more ATP per amount of resource consumed, but may come at the cost of increased reactive oxygen species (ROS) production damaging tissues through oxidative stress. Mitochondrial efficiency is a plastic trait but how it changes through postnatal development remains unknown. We hypothesized that strong selective pressure could lead to an increased mitochondrial efficiency to support fast growth but incur an oxidative cost. We tested this hypothesis by quantifying mitochondrial aerobic metabolism, efficiency and ROS production through postnatal growth in Japanese quail (*Coturnix japonica*), in two highly aerobic tissues: skeletal and cardiac muscles. Mitochondrial efficiency was indeed higher during peak growth in both tissues, but this was surprisingly associated with markedly lower ROS production. This high efficiency was likely achieved via both a lower proton leak and a higher contribution of complex I to respiration. These results show that enhancing mitochondrial efficiency may be important to support growth, but suggest the presence of unexpected ROS mitigation processes during early-life growth.

## Introduction

Early-life growth rates are under strong selective pressures (Arendt, 1997; Remeŝ & Martin, 2002). In many species, increased growth rate and body mass are associated with higher survival and reproductive success (Stearns & Koella, 1986). However, animals obtain energy for growth from food through foraging, which is costly (Biro et al., 2006; Dmitriew, 2011). Yet, energy available for the animal does not only depend on food uptake, but also on its processing. In this context, the conversion of food-derived molecules to a form of energy available at the cellular level (i.e. adenosine tri-phosphate, ATP) is mainly driven by mitochondrial aerobic metabolism (Koch et al., 2021). Mitochondrial electron transport system (ETS) uses electrons harvested from food-derived molecules and O_2_ as a final electron acceptor to drive ATP synthesis, but the efficiency at which ATP is produced is variable (Salin et al., 2015). Mitochondrial efficiency depends on 1. the amount of protons leaking from the intermembrane space back to the mitochondrial matrix without being coupled to ATP synthesis, passively due to membrane composition and high membrane potential, or actively through specific transporters (Jastroch et al., 2010); 2. the type of food-derived substrate being used, with substrates providing electrons entry through complex II leading to less ATP being synthesized by units of O_2_ consumed than those entering through complex I (Brand, 2005). Thus, mitochondrial efficiency can be defined as the amount of ATP produced per atom of oxygen consumed (ATP/O), or to a lesser extent by the proportion of O_2_ consumed that is coupled to ATP production (Salin et al., 2018).

Mitochondrial efficiency is a plastic trait, as it can change quickly in response to fasting (Roussel et al., 2018) or for reproduction (Hood, 2024). Yet, consistent inter-individual differences in mitochondrial efficiency have also been demonstrated (Stier et al., 2019, 2022), which contributes to suggest that variation in mitochondrial efficiency may underlie variation in animal performance (Koch et al., 2021; Salin et al., 2015). With efficient mitochondria, more energy can be taken from food, but high efficiency may lead to increased reactive oxygen species (ROS) production (Berry et al., 2018). Overproduction of ROS that exceeds cell detoxification capacities can cause oxidative damage, threatening cell integrity and survival (Speakman et al., 2015). On the contrary, slightly increasing proton leak (and thus reducing mitochondrial efficiency) seems to reduce ROS production (Brand, 2000). This highlights a potential trade-off between the efficiency of energy transduction and cell protection (Hood et al., 2018; Koch et al., 2021).

Inter-individual variation in growth performances can partly be explained by inter-individual variation in mitochondrial efficiency. Indeed, growth rate is positively associated with mitochondrial efficiency in fish (Salin et al., 2019), domestic chicken (broiler versus layers in (Sirsat & Dzialowski, 2018)) and frogs (Salin, Roussel, et al., 2012). Yet, fast growth has been associated with high ROS production and oxidative stress, which themselves can contribute to decrease growth rate through an oxidative constraint (Smith et al., 2016). Growth rate could thus be limited by ROS production, but an experimental manipulation decreasing mitochondrial efficiency in frog tadpoles led to lower ROS production and slowed-down development (Salin, Luquet, et al., 2012). Thus, the trade-off between mitochondrial efficiency and ROS production seems to play a role in growth rate determination (Koch et al., 2021; Monaghan et al., 2009). Yet, the changes in mitochondrial efficiency and ROS production throughout ontogeny remain mostly unknown.

To fill this gap, we studied mitochondrial aerobic metabolism, efficiency and ROS production during early-life in Japanese quails, using two highly aerobic organs, namely heart and skeletal muscles. We hypothesized that quails would optimize their metabolism during the early-life fast growth resulting in a higher mitochondrial efficiency in young versus adult quails, but at the cost of higher ROS production.

## Material and methods

### 1. General procedures

Japanese quail (*Coturnix japonica*) from two artificial selection lines (E+: high fear response; E-: low fear response; Faure et al. 2006) were obtained from INRAE Nouzilly (France) at an age of 7 days post-hatching, and maintained in one indoor aviary at *ca*. 22°C on a 12 L : 12 D cycle with water and food *ad libitum*. The artificial selection lines were the focus of another study, and thus the identity of the selection line E+ vs. E-is not included in the analyses. Fourteen quails (7 E+, 7 E-; 9 females, 5 males) were used for monitoring body mass growth from day 10 (early-growth) to day 60 (sexual maturity), with regular weighing (Fig. 1). Twenty-nine other quails (14 E+, 15 E-; 14 females, 15 males) were used for mitochondrial analyses, and were euthanized by cervical dislocation in three groups of age: day 16 (16.0 ± 1.3 days) corresponding to early peak growth (Fig. 1), day 25 (25.0 ± 4.3 days) corresponding to late peak growth (Fig. 1) and day 60 (60.0 ± 3.0 days) corresponding to post-growth and sexually mature individuals. Following euthanasia, the heart and one piece of pectoral skeletal muscle were immediately sampled and placed in an ice-cold mitochondrial isolation buffer (100 mM sucrose, 50 mM KCl, 5 mM EDTA and 50 mM Tris-base, pH 7.4).

**Fig. 1:**
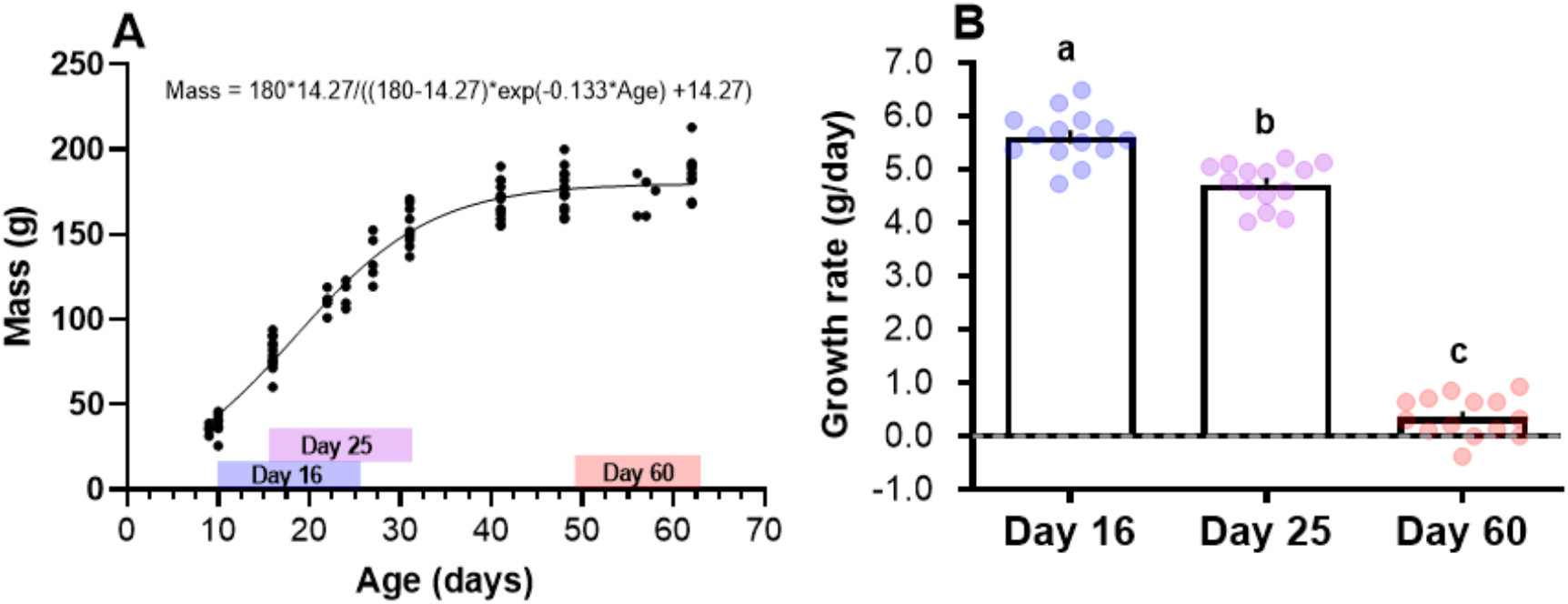
Body mass growth of Japanese quails. A) Logistic growth curve based on 14 individuals repeatedly weighed during growth (N = 14, n = 98). Colored rectangles correspond to the time period used to calculate daily growth rates for panel B. B) Growth rates around the 3 age points at which birds were sampled for mitochondrial analyses. Letters indicate significant differences (p ≤ 0.050) according to post-hoc tests with Tukey correction.

### 2. Mitochondria isolation

Tissue samples were finely cut up, homogenized using a Potter–Elvehjem homogenizer (5 passages), and centrifuged at 1000 g for 10 min (4°C). The supernatant with subsarcolemmal mitochondria was conserved in an ice bath, and the pellet with intermyofibrillar mitochondria was suspended in isolation buffer and digested with protease from *Bacillus licheniformis* (1 mg.g^−1^ of muscle) for 5 min in an ice bath. The homogenate was then diluted (1:2) using the supernatant from the first centrifugation, and the mixture was centrifuged during 10 min at 1000 g. The resulting supernatant was filtered and centrifuged during 10 min at 8700 g to pellet mitochondria. The mitochondrial pellet was washed twice by resuspension in isolation buffer and centrifugation at 8700 g for 10 min. Quantification of mitochondrial proteins in the suspension were performed using the Biuret method, with bovine serum albumin as standard.

### 3. Mitochondrial respiration and ATP production

The rates of mitochondrial respiration and ATP synthesis were assessed at 40°C as described previously (Barbe et al., 2023). The rate of oxygen consumption (Clark oxygen electrode, Rank Brothers Ltd, UK) was recorded in respiratory buffer for mitochondria (120 mM KCl, 5 mM KH2PO4, 2 mM MgCl2, 1 mM EGTA, 3 mM HEPES and 0.30% BSA, pH 7.4), supplemented with an ADP-regenerating system consisting of 20 mM glucose and 2 U.ml^−1^ hexokinase. Respiration was initiated by adding complex I substrates 5 mM pyruvate and 2.5 mM malate (*LEAK*_*CI*_ respiration with complex I substrates), followed by the addition of complex II substrate 5 mM succinate (*LEAK*_*CI+II*_ respiration, with complex I and II substrates). Then, 500μM ADP was added to stimulate oxidative phosphorylation (*OXPHOS*_*CI+II*_ respiration). The mitochondrial OXPHOS control efficiency (OxCE) was calculated as (OXPHOS_CI+II_ -LEAK_CI+II_)/OXPHOS_CI+II_ (Gnaiger, 2020), and the relative contribution of electron entry through complex I and complex II as (LEAK_CI+II_ -LEAK_CI_)/LEAK_CI+II_.

After oxygen consumption was monitored, four aliquots of mitochondrial suspension were withdrawn every minute and quenched in a perchloric acid solution (10% HClO4, 25 mM EDTA). The denatured proteins were centrifuged at 21,000 g for 5 min at 4°C. The supernatant was neutralized with a KOH solution (2 M KOH, 0.3 M Mops) and centrifuged at 21,000 g for 5 min (4°C). The resulting supernatant was incubated at room temperature in reaction buffer (50 mM triethanolamine-HCl, 7.5 Mm MgCl2 and 3.75 mM EDTA, pH 7.4), supplemented with 0.5 mM NAD and assayed spectrophotometrically at 340 nm to monitor the NADH production in the presence of 0.5 U glucose-6-phosphate dehydrogenase (from *Leuconostoc mesenteroides*). ATP production rate was thus calculated from the slope of glucose-6-phosphate accumulation over the sampling time intervals. ATP synthesis was also determined in a basal non-phosphorylating respiration rate in the presence of 2 μg ml^−1^ oligomycin to ensure that the measured rates were specific to the mitochondrial ATP synthase activity. These values were taken into account to calculate the rate of mitochondrial ATP synthesis. From ATP production rate, the ATP/O ratio was determined by dividing ATP production by the corresponding OXPHOS respiration.

### 4. Mitochondrial reactive oxygen species production

The rate of H_2_O_2_ released by skeletal muscle and heart mitochondria was measured in respiratory buffer supplemented with 5 U.mL horseradish peroxidase and 1 μM Amplex® Red fluorescent dye at 40°C using a 96-well fluorescence plate reader (Fluorescence spectrophotometer F-7100, Hitachi) at excitation and emission wavelengths of 560 and 584 nm, respectively. The kinetics of H_2_O_2_ production were monitored for 10 minutes, and the slope of fluorescence accumulation was converted to H_2_O_2_ values (pmol.min-1.mg protein-1) using an H_2_O_2_ standard curve ranging from 4.5 to 36 pmol. The rate of H_2_O_2_ was assessed under both OXPHOS_*CI+II*_ and LEAK_*CI+II*_ conditions, as for respiration rates above. Coefficients of variation based on duplicate measurements were low: 7.7 ± 1.1%.

### 5. Statistical analyses

Body mass data was fitted with a logistic growth curve (Fig. 1A), corresponding to Y=YM*Y0/((YM-Y0)*exp(-k*x) +Y0), with *Y* being the body mass (g) at a given age, *x* the age (days), *Y0* the starting body mass, *YM* the asymptotic body mass, *k* the growth rate constant and 1/k the age of the first inflection point. Growth rate (in g.day^-1^) around the sampling times for mitochondrial analyses (day 16, 25 and 60) were calculated from body mass data around those ages, and analyzed using a linear mixed model including Age (day 16 vs. day 25 vs. day 60), Line (E+ vs. E-), Sex (Male vs. Female) and their two-ways interactions as fixed effects, as well as bird identity as a random intercept to account for repeated measurements over time. Free radical electron leak (ROS/O ratio) was calculated as the fraction (%) of oxygen consumption that is reduced into H_2_O_2_ at the respiratory chain instead of H_2_O at the cytochrome-c oxidase (Roussel & Voituron, 2020). The oxidative cost of mitochondrial ATP synthesis was calculated from the ratio of H_2_O_2_ generation under phosphorylating state divided by the corresponding rate of ATP synthesis (ROS/ATP ratio; (Roussel & Voituron, 2020)). All mitochondrial parameters were analyzed using linear mixed models including Tissue (skeletal vs. cardiac muscles), Age, and their two-way interaction as fixed effects, as well as bird identity as a random intercept to account for repeated measurements (i.e. two tissues of the same birds). Sex and selection lines were initially included in statistical analyses, but were never significant predictors of mitochondrial traits as main effects. They were thus removed from final models.

All analyses were performed using R version 4.4.2, using the *lme4* (Bates et al., 2015) and *LmerTest* (Kuznetsova et al., 2017) packages for mixed models, and the *emmeans* package (Lenth et al., 2019 for post-hoc tests with Tukey adjustment. Distributional assumptions were checked using the *DHARMa* package (Hartig & Hartig 2017). Two minor violations of assumptions occurred but linear mixed models are robust to such minor violations (Schielzeth et al., 2020). Means are always presented ± SE.

## Results

### 1. Growth

Japanese quails exhibit a logistic growth curve typical of most avian models (Fig. 1A). Body mass growth was significantly influenced by age (*F*_*2,26*_ = 683.9, p < 0.001), being higher around day 16 than later on, and significantly higher at day 25 than at day 60 (Fig. 1B).

### 2. Mitochondrial respiration and ATP production rates

*LEAK*_*CI+II*_ respiration was marginally affected by the interaction between Tissue and Age (Fig. 2A; *F*_*2,26*_ = 2.8, p = 0.081). Specifically, *LEAK*_*CI+II*_ only differed between tissues at day 60, being higher in cardiac than skeletal muscle (Fig. 2A; *t*_26_ = 3.7, p = 0.001). In skeletal muscle, *LEAK*_*CI+II*_ was higher at day 25 than day 16 and 60 (Fig. 2A; day 16 vs. 25: *t*_51.3_ = -2.7, p = 0.023; day 25 vs. 60: *t*_51.3_ = 2.4, p = 0.050). The pattern was different in cardiac muscle, with *LEAK*_*CI+II*_ increasing from day 16 onward (Fig. 2A; day 16 vs. 25: *t*_51.3_ = -2.7, p = 0.027; day 16 vs. 60: *t*_51.3_ = -3.0, p = 0.011).

**Fig. 2:**
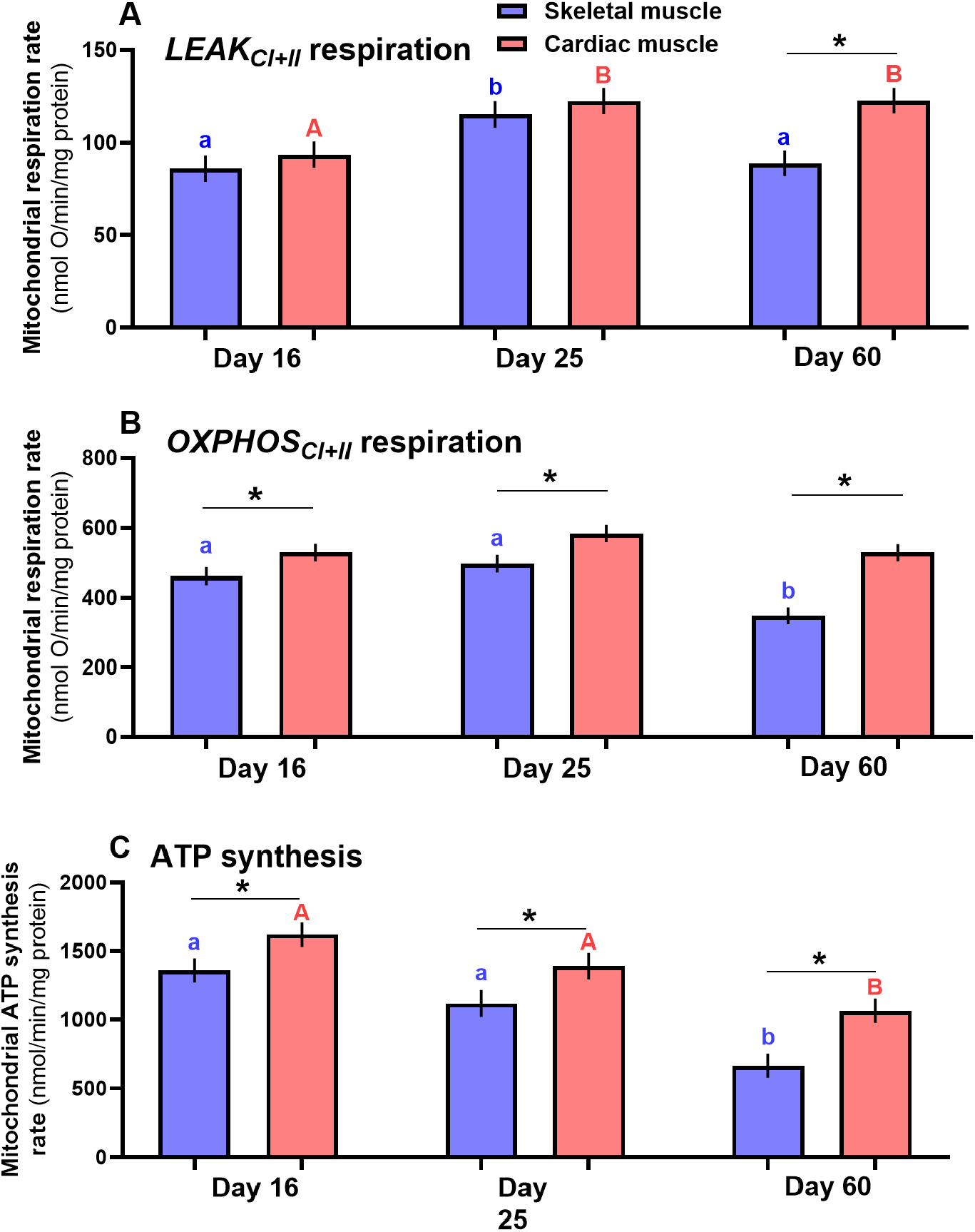
Age-related changes in mitochondrial respiration and ATP production rates in skeletal and cardiac muscles of Japanese quails. A) Mitochondrial *LEAK*_*CI+II*_ respiration, corresponding to O_2_ consumed by mitochondria while no ATP synthesis occurs. B) Mitochondrial *OXPHOS*_*CI+II*_ respiration, corresponding to O_2_ consumed by mitochondria during active ATP synthesis. C) Mitochondrial ATP production. Letters indicate significant age differences within tissue, and * significant differences between tissues for a given age (p ≤ 0.050) according to post-hoc tests with Tukey correction (N = 29, n = 58).

*OXPHOS*_*CI+II*_ respiration was significantly affected by the interaction between Tissue and Age (Fig. 2B; *F*_*2,26*_ = 4.6, p = 0.019). *OXPHOS*_*CI+II*_ was consistently higher in cardiac than in skeletal muscle (Fig. 2B; all *t*_26_ > 2.2 and all p < 0.032), and was not influenced by Age in cardiac muscle (all *t*_47.5_ < 1.7 and all p > 0.23). Conversely, OXPHOS_*CI+II*_ significantly decreased from active growth (*i*.*e*. days 16 and 25) to adulthood at day 60 in skeletal muscle (Fig. 2B; day 16 vs. 60: *t*_47.5_ = 3.3, p = 0.005; day 25 vs. 60: *t*_47.5_ = 4.4, p < 0.001).

ATP synthesis rate was significantly higher in cardiac than skeletal muscle (*F*_*1,26*_ = 29.3, p < 0.001), and decreased with age consistently in both tissues (Fig. 2C; *F*_*2,26*_ = 24.1, p < 0.001), being higher during active growth than at adulthood (day 16 vs. 60: *t*_26_ = 6.7, p < 0.001; day 25 vs. 60: *t*_26_ = 4.9, p < 0.001). The interaction between Age and Tissue was however not significant (*F*_*2,26*_ = 0.6, p = 0.56).

### 3. Mitochondrial efficiency and complex II contribution to respiration

*OXPHOS* control efficiency was significantly higher in cardiac than skeletal muscle (*F*_*1,26*_ = 5.2, p = 0.031), and decreased progressively with age (Fig. 3A; *F*_*2,26*_ = 15.3, p < 0.001; day 16 vs. 25: *t*_26_ = 3.1, p = 0.014; day 16 vs. 60: *t*_26_ = 5.5, p < 0.001; day 25 vs. 60: *t*_26_ = 2.5, p = 0.044). Mitochondrial efficiency measured as ATP/O ratio also decreased progressively with Age (Fig. 3B; *F*_*2,26*_ = 20.1, p < 0.001, day 16 vs. 25: *t*_26_ = 3.6, p = 0.004; day 16 vs. 60: *t*_26_ = 6.3, p < 0.001; day 25 vs. 60: *t*_26_ = 2.8, p = 0.023), but did not differ between tissues, neither as a main factor (*F*_*1,26*_ = 1.9, p = 0.18), nor in interaction with age (*F*_*2,26*_ = 0.01 p = 0.99).

**Fig. 3:**
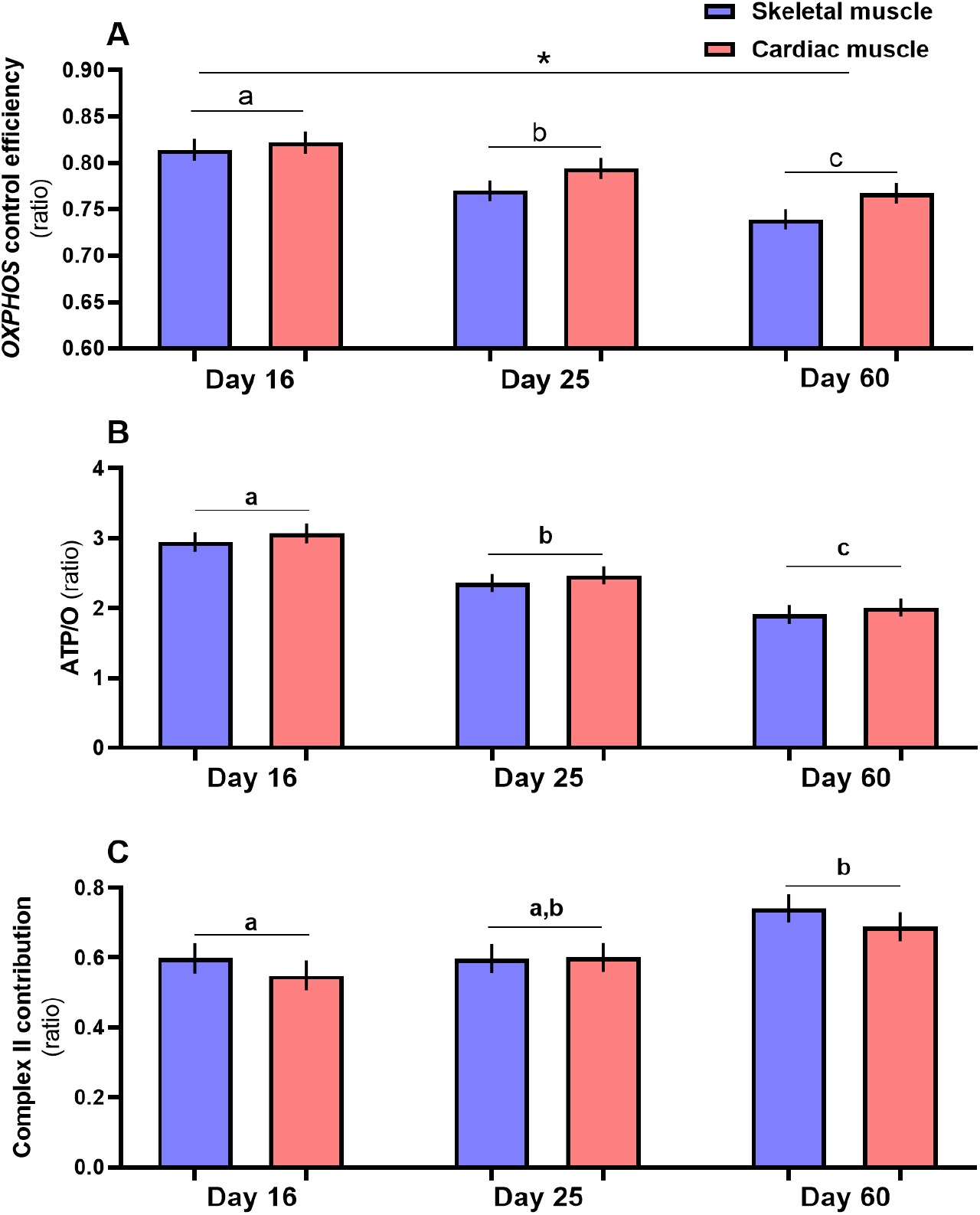
Age-related changes in mitochondrial efficiency and substrate use in skeletal and cardiac muscles of Japanese quails. A) *OXPHOS* control efficiency corresponds to the proportion of mitochondrial O_2_ consumed devoted to ATP synthesis. B) Mitochondrial coupling efficiency (ATP/O ratio) corresponds to the amount of ATP molecule produced per atom of oxygen consumed, C) Complex II contribution corresponds to the proportion of electron entry through complex II when both complex I and II are fueled simultaneously to mitochondria. Letters indicate significant age differences, and * significant differences between tissues (p ≤ 0.050) according to post-hoc tests with Tukey correction (N = 29, n = 58).

The relative contribution of complex II to respiration progressively increased with age (Fig. 3C; *F*_*2,26*_ = 4.0, p = 0.031; day 16 vs. 25: *t*_26_ = -0.5, p = 0.88; day 16 vs. 60: *t*_26_ = -2.6, p = 0.037; day 25 vs. 60: *t*_26_ = -2.2, p = 0.089) but was not significantly influenced by Tissue (*F*_*1,26*_ = 2.7, p = 0.11) nor by the Tissue*Age interaction (*F*_*2,26*_ = 0.9, p = 0.41).

### 4. Mitochondrial ROS production

ROS production under a non-phosphorylating state (i.e. *LEAK*_*CI+II*_) measured as H_2_O_2_ efflux was significantly higher in adulthood than during growth (Fig. 4A, *F*_*2,26*_ = 30.8, p < 0.001; day 16 vs. 25: *t*_26_ = 2.0, p = 0.14; day 16 vs. 60: *t*_26_ = -5.4, p < 0.001; day 25 vs. 60: *t*_26_ = -7.6, p < 0.001), but was not significantly influenced by Tissue (*F*_*1,26*_ = 0.1, p = 0.81) nor by the Age*Tissue interaction (*F*_*2,26*_ = 2.2, p = 0.14). During active ATP synthesis (i.e. *OXPHOS*_*CI+II*_), the pattern was slightly different, with H_2_O_2_ efflux being significantly impacted by the Age*Tissue interaction (Fig. 4B; *F*_*2,26*_ = 3.4, p = 0.049). While ROS production was higher in both tissues at adulthood than during active growth (day 16 and 25 vs. day 60: all *t*_34.1_ < -5.5, p < 0.001), cardiac mitochondria produced significantly more ROS than skeletal muscle mitochondria at day 16 (Fig. 4B; *t*_26_ = 5.1, p < 0.001) and day 60 (*t*_26_ = 2.5, p = 0.019), but not significantly so at day 25 (*t*_26_ = 1.8, p = 0.089)

**Fig. 4:**
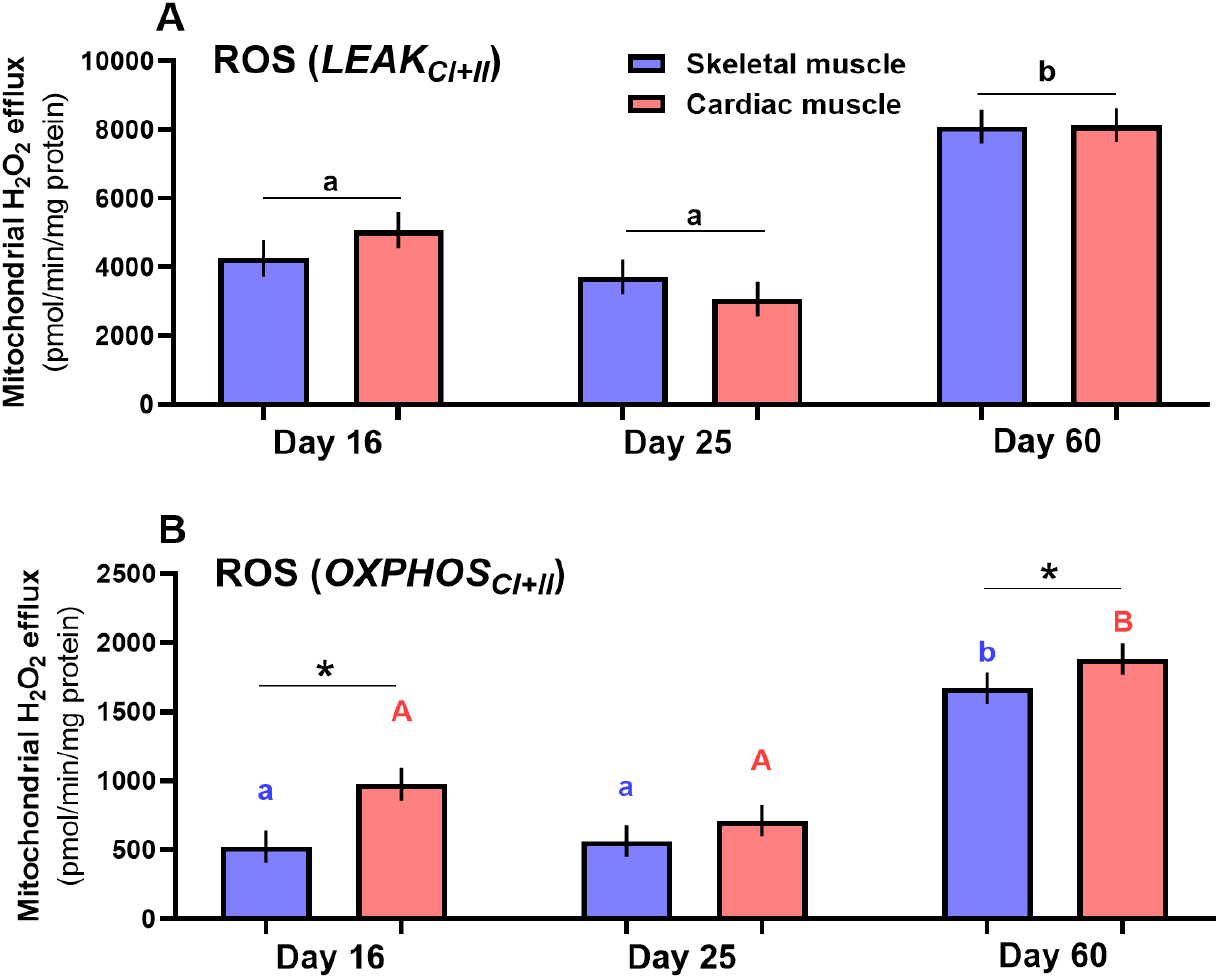
Age-related changes in mitochondrial ROS production (H_2_O_2_ efflux) in skeletal and cardiac muscles of Japanese quails. A) H_2_O_2_ efflux during *LEAK*_*CI+II*_ state, where ROS production is high. B) H_2_O_2_ efflux during *OXPHOS*_*CI+II*_ state, where ROS production is limited by active electron transport driven by ATP synthesis. Letters indicate significant age differences within tissue, and * significant differences between tissues for a given age (p ≤ 0.050) according to post-hoc tests with Tukey correction (N = 29, n = 58).

### 5. Electron leak

Electron leak under a non-phosphorylating state (i.e. *LEAK*_*CI+II*_) was significantly impacted by the Age*Tissue interaction (*F*_*2,26*_ = 4.2, p = 0.026; Fig. 5A). In skeletal muscle, it was higher at adulthood than during active growth (Fig. 5A; day 16 vs. 25: *t*_46_ = 1.7, p = 0.20;day 16 vs. 60: *t*_46_ = -4.3, p < 0.001; day 25 vs. 60: *t*_46_ = -6.1, p < 0.001), while in cardiac muscle it slightly decreased from day 16 to 25 (*t*_46_ = 3.2, p = 0.006) but then increased at adulthood (day 25 vs. day 60: *t*_46_ = -4.4, p < 0.001). Skeletal muscle mitochondria had more electrons leaking in the *LEAK* stage than cardiac muscle mitochondria, but only at adulthood (Fig. 5A; day 16: *t*_26_ = 0.8, p = 0.42; day 25: *t*_26_ = -1.1, p = 0.30; day 60: *t*_26_ = -3.3, p = 0.003).

**Fig. 5:**
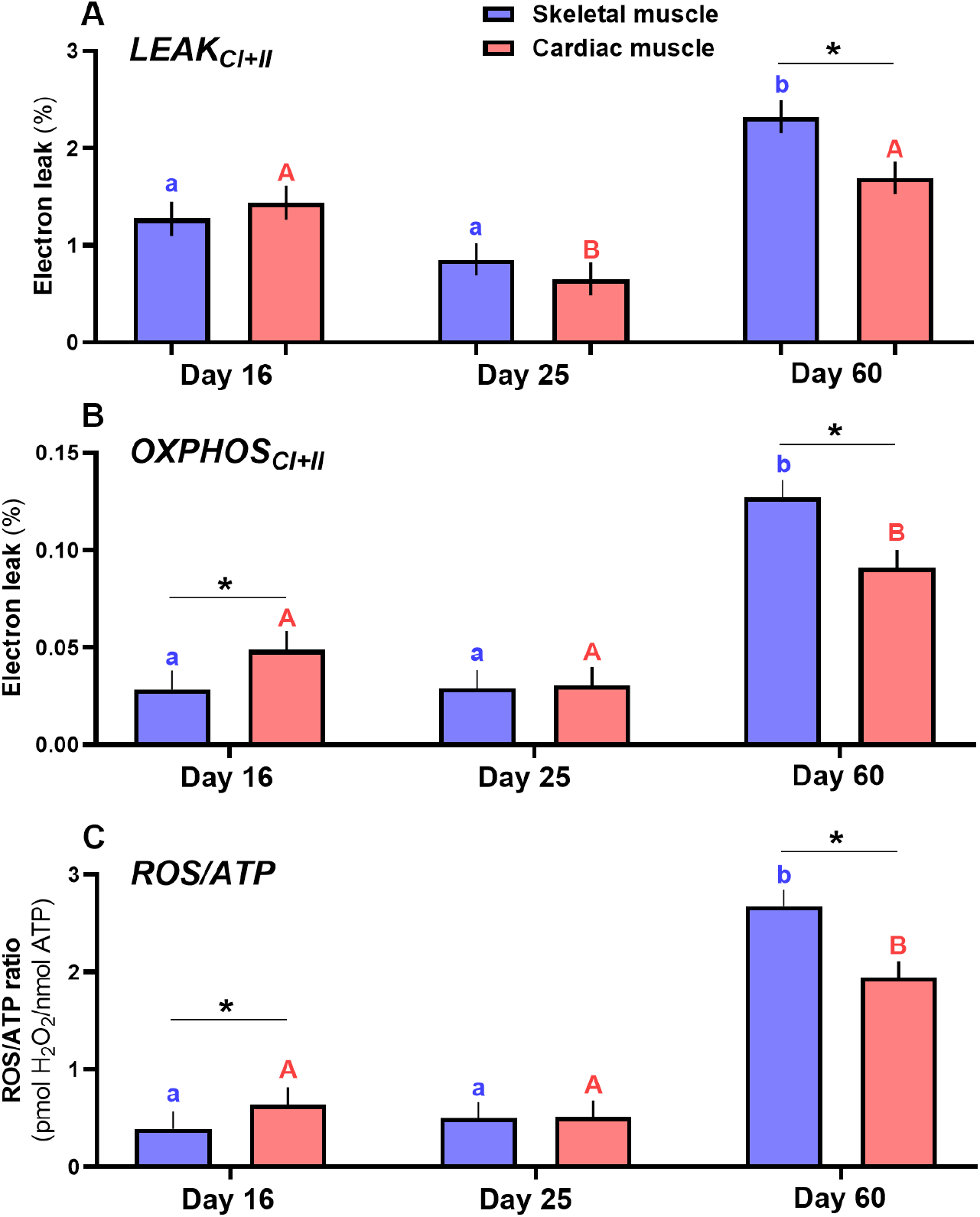
Age-related changes in mitochondrial electron leak (A-B) and oxidative cost of ATP production(C) in skeletal and cardiac muscles of Japanese quails. Electron leak corresponds to the percentage of electrons escaping the electron transport system under A) *LEAK* conditions, (B) *OXPHOS* conditions. The oxidative cost of ATP production is calculated as the amount of ROS produced per unit of ATP produced. Letters indicate significant age differences within tissue, and * significant differences between tissues for a given age (p ≤ 0.050) according to post-hoc tests with Tukey correction (N = 29, n = 58).

Under phosphorylating conditions (i.e. *OXPHOS*_*CI+II*_), electron leak was also modulated by the Age*Tissue interaction but in a slightly different fashion (*F*_*2,26*_ = 15.3, p < 0.001; Fig. 5B). In both tissues, electron leak in the *OXPHOS* stage was more pronounced at adulthood than during active growth (day 16 and 25 vs. day 60: all *t*_35.2_ < -3.1 and all p < 0.009), and more pronounced in cardiac muscle during early-growth (day 16: *t*_26_ = 2.7, p = 0.013) but in the skeletal muscle at adulthood (*t*_26_ = -5.0, p < 0.001)

### 6. Oxidative cost of ATP production

The amount of ROS produced per unit of ATP synthesized was significantly influenced by the Age*Tissue interaction (*F*_*2,26*_ = 16.4, p < 0.001; Fig. 5C). Specifically, ROS/ATP was higher at adulthood than during active growth in both tissues (day 16 and 25 vs. day 60: all *t*_34.2_ < - 5.3 and all p < 0.001), and was higher in skeletal than cardiac muscle, but only at adulthood (Fig. 5C; day 16: *t*_34.2_ = 1.9, p = 0.072; day 25: *t*_34.2_ = 0.1, p = 0.92; day 60: *t*_34.2_ = -5.9, p < 0.001).

## Discussion

Owing to oxygen, ATP and H_2_O_2_ monitoring of isolated mitochondria from two aerobic tissues (skeletal and cardiac muscles) of Japanese quails, we showed that mitochondrial efficiency is higher during the active growth period compared to adulthood. Contrary to expectations, H_2_O_2_ levels were higher in adults than during active growth, resulting in a higher oxidative cost of ATP production in adults than in growing individuals.

Mitochondrial efficiency, measured both as OXPHOS control efficiency and ATP/O ratio, was higher during early growth than at adulthood. While both kind of indicators of efficiency have been suggested to convey different or even sometimes opposite results (Salin et al., 2018), our results suggest that the increased mitochondrial efficiency observed during growth is robust, and also identical between two aerobic tissues, despite widespread tissue-specific effects on mitochondrial responses (e.g. Chausse et al. (2015)). Interestingly, previous studies showed that increased mitochondrial efficiency is associated with increased higher growth performances (Salin et al., 2019; Salin, Roussel, et al., 2012; Sirsat & Dzialowski, 2018). Here, we showed that mitochondrial efficiency is increased during growth compared to adulthood. These results are ecologically relevant, suggesting that growing quails optimize their mitochondrial efficiency to minimize the cost of ATP production, as the OXPHOS control efficiency and ATP/O ratios determine how much nutrients and oxygen are needed to match energy demand. Therefore, it seems that mitochondrial plasticity is at play during growth with mitochondrial efficiency being optimized to maximize energy availability from limited resources during an energy-demanding period.

The age-related variation in mitochondrial efficiency we observed may be linked to two main processes: 1. the redox quotient, i.e. NADH/FADH_2_ ratio, that drives how many electrons enter complex I relative to complex II (Brand, 2005) ; 2. Proton leak reactions that negatively impact mitochondrial efficiency (Brand et al., 1994). Although the redox quotient was not directly quantified in the present study, we found an increased succinate contribution ratio over the course of quail development, indicating that muscle mitochondria from quail may rely more on FADH_2_-linked substrate at adulthood than in early life. This slight shift toward complex II supported respiration may contribute to lower coupling efficiency in adult quails. The type of substrate being oxidized also affects mitochondrial efficiency. For instance, fatty acid oxidation leads to a lower mitochondrial efficiency than carbohydrate oxidation because it provides more electron entry through FADH_2_-linked complexes (Brand, 2005). Birds are well known to maintain high lipid oxidation throughout life and during high demanding energetic challenges (Barbe et al., 2023; Vaillancourt et al., 2005). Although mitochondrial substrates used during growth and adulthood are not known to differ, the lower succinate contribution ratio reported in fast growing quails would favor a channeling of energy from electron transport toward ATP synthesis and growth performance.

Mitochondrial efficiency is known to be affected by the amount of protons leaking from the intermembrane space to the matrix. Based on our results from OXPHOS vs. LEAK respiration rates (and thus OXPHOS control efficiency), the age-related variation in mitochondrial efficiency is linked to a lower proton leakage during growth. Yet, the source of the reduced proton leakage during growth remains to be determined, as proton leak can occur passively due to membrane composition (i.e. proportion of polyunsaturated fatty acids in the inner mitochondrial membrane) and high membrane potential, or actively through specific transporters, *e*.*g*. ANT and avUCP. Investigating the lipid composition of mitochondrial membranes and the expression of specific proteins may help to clarify the potential involvement of these two pathways (Brand et al., 1994; Kikusato & Toyomizu, 2013).

Many aspects of mitochondrial metabolism are also known to negatively scale with body mass at the inter-specific level, which include proton leak activity, inner membrane proton conductance, OXPHOS and ATP synthesis rates (Barbe et al., 2023; Boël et al., 2019, 2023; Brand et al., 2003; Porter & Brand, 1993). Yet, the maximal coupling efficiency of liver, skeletal muscle and heart mitochondria, calculated as the ratio between ATP synthesis rate and corresponding OXPHOS respiration rate, were independent of body mass (Barbe et al., 2023; Boël et al., 2019, 2023). In bird species ranging from 15 g to 150 kg (Barbe et al., 2023), the values of mitochondrial coupling efficiency average 2.06 ± 0.07 and 2.18 ± 0.08 in skeletal muscle and heart mitochondria, respectively, which are similar to the ATP/O ratios calculated in the present study for adult quail muscle mitochondria. Therefore, the higher coupling efficiency of active growing quails is unlikely to be explained by body mass difference with adult individuals.

Despite a higher mitochondrial efficiency, we found lower H_2_O_2_ production, oxidative cost of ATP (%H_2_O_2_/ATP), and OXPHOS electron leak (fraction of total electron flow reducing O_2_ into oxygen free radicals instead of O_2_ into H_2_O at complex IV) in early life compared to adulthood. This is contrary to our initial prediction since increased mitochondrial efficiency has been linked theoretically and empirically to increased ROS production (Brand, 2000; Korshunov et al., 1997). Indeed, proton leak has been suggested to play a role in ROS production regulation since increased proton leak (and thus decreased mitochondrial efficiency) is expected to reduce the proton motive force, resulting in a lower probability of electron leak (Berry et al., 2018). However, young quails have a high ATP demand due to growth that could explain why they are able to increase their mitochondrial efficiency without having increased ROS production compared to adults. Indeed, high ATP demand increases proton flux through the ATP synthase, resulting in a decrease of proton motive force, and thus a lower likelihood of electron leak without a need for proton leak and loss of efficiency. The phosphorylating condition of young quail mitochondria could therefore be a major factor influencing ROS production (Quinlan et al., 2012). In addition, the lower succinate contribution to respiration in fast growing quails may drive a lower proton motive force (Mookerjee et al., 2021), which would also participate in lowering mitochondrial H_2_O_2_ release compared to adulthood. Altogether, a high ATP demand and a low succinate contribution ratio could explain how young quails are able to maintain both high mitochondrial efficiency and low ROS production whereas high mitochondrial efficiency is generally linked to increased ROS in static conditions. An alternative explanation to the reduced mitochondrial H_2_O_2_ efflux we observed during growth may also be linked to differences in intra-mitochondrial ROS scavenging capacity (e.g. linked to superoxide dismutase SOD2 isoform) of mitochondria since ROS production takes place into the matrix and we measured H_2_O_2_ release from the mitochondria (Munro & Treberg, 2017; Quinlan et al., 2012). If a higher ROS production and ROS/ATP ratio in adulthood would theoretically impair cellular integrity (López-Lluch et al., 2006; Neretti et al., 2009), there could be compensation through increased endogenous antioxidant defenses and repair processes. Indeed, adult quails were not found to be more affected by oxidative damage than growing ones, when measured as DNA oxidative damage in blood cells (Stier et al., 2020). A proper assessment of antioxidants and oxidative damage across tissues will be needed to test this hypothesis.

While the classical physiological trade-off between mitochondrial efficiency and ROS production has been suggested to underlie the life-history trade-off between growth and longevity, our results suggest this is not the case. Plasticity in mitochondrial metabolism enables to reach both high efficiency and low ROS production, thus opening up the question of why this apparent optimal functioning is not maintained at adulthood.

## Author contribution

AS designed the study, with input from DR and LC. AS, AC and LA conducted the experimental procedures on birds. AS, DR and JB conducted laboratory work. LL and AS analyzed the data and wrote the manuscript, with input from all other authors.

## Acknowledgements

We are grateful to Carsten Schradin and Lindelani Makuya for the scientific writing course they organized in which LL started to write this manuscript.

## Data accessibility

All data and R scripts are available in FigShare (10.6084/m9.figshare.31939719).

## Ethical permits

All procedures were approved by the French Government under the reference APAFIS #34983-2022012511537353 v4, following positive evaluation by the Comité d’Expérimentation Animale de l’Université Claude Bernard Lyon 1 – CEEA-55.

## References

Arendt, J. D. (1997). Adaptive Intrinsic Growth Rates : An Integration Across Taxa. The Quarterly Review of Biology, 72(2), 149–177. 10.1086/419764

Barbe, J., Watson, J., Roussel, D., & Voituron, Y. (2023). The allometry of mitochondrial efficiency is tissue dependent:A comparison between skeletal and cardiac muscles of birds. Journal of Experimental Biology, 226(23), jeb246299. 10.1242/jeb.246299

Bates, D., Mächler, M., Bolker, B., & Walker, S. (2015). Fitting Linear Mixed-Effects Models Using lme4. Journal of Statistical Software, 67, 1–48. 10.18637/jss.v067.i01

Berry, B. J., Trewin, A. J., Amitrano, A. M., Kim, M., & Wojtovich, A. P. (2018). Use the Protonmotive Force:Mitochondrial Uncoupling and Reactive Oxygen Species. Journal of Molecular Biology, 430(21), 3873–3891. 10.1016/j.jmb.2018.03.025

Biro, P. A., Abrahams, M. V., Post, J. R., & Parkinson, E. A. (2006). Behavioural trade-offs between growth and mortality explain evolution of submaximal growth rates. Journal of Animal Ecology, 75(5), 1165–1171. 10.1111/j.1365-2656.2006.01137.x

Boël, M., Romestaing, C., Voituron, Y., & Roussel, D. (2019). Allometry of mitochondrial efficiency is set by metabolic intensity. Proceedings of the Royal Society B: Biological Sciences, 286(1911), 20191693. 10.1098/rspb.2019.1693

Boël, M., Voituron, Y., & Roussel, D. (2023). Body mass dependence of oxidative phosphorylation efficiency in liver mitochondria from mammals. Comparative Biochemistry and Physiology Part A, Molecular & Integrative Physiology, 284, 111490. 10.1016/j.cbpa.2023.111490

Brand, M. D. (2000). Uncoupling to survive ? The role of mitochondrial inefficiency in ageing. Experimental Gerontology, Biological Aging-Euroconference on Molecular, Cellular and Tissue Gerontology, 35(6), 811–820. 10.1016/S0531-5565(00)00135-2

Brand, M. D. (2005). The efficiency and plasticity of mitochondrial energy transduction. Biochemical Society Transactions, 33(5), 897–904. 10.1042/BST0330897

Brand, M. D., Chien, L.-F., Ainscow, E. K., Rolfe, D. F. S., & Porter, R. K. (1994). The causes and functions of mitochondrial proton leak. Biochimica et Biophysica Acta (BBA)-Bioenergetics, 1187(2), 132–139. 10.1016/0005-2728(94)90099-X

Brand, M. D., Turner, N., Ocloo, A., Else, P. L., & Hulbert, A. J. (2003). Proton conductance and fatty acyl composition of liver mitochondria correlates with body mass in birds. Biochemical Journal, 376(3), 741–748. 10.1042/bj20030984

Chausse, B., Vieira-Lara, M. A., Sanchez, A. B., Medeiros, M. H. G., & Kowaltowski, A. J. (2015). Intermittent Fasting Results in Tissue-Specific Changes in Bioenergetics and Redox State. PLOS ONE, 10(3), e0120413. 10.1371/journal.pone.0120413

Dmitriew, C. M. (2011). The evolution of growth trajectories:What limits growth rate? Biological Reviews, 86(1), 97–116. 10.1111/j.1469-185X.2010.00136.x

Gnaiger, E. (2020). Mitochondrial pathways and respiratory control:An Introduction to OXPHOS Analysis. 5th ed. Bioenergetics Communications, 2020, 2–2. 10.26124/bec:2020-0002

Hartig, F. and Hartig, M.F., 2017. Package ‘dharma’. R package, 531(532), p.435.

Hood, W. R. (2024). A Mitochondrial Perspective on the Demands of Reproduction. Integrative and Comparative Biology, 64(6), 1611–1622. 10.1093/icb/icae049

Hood, W. R., Zhang, Y., Mowry, A. V., Hyatt, H. W., & Kavazis, A. N. (2018). Life History Trade-offs within the Context of Mitochondrial Hormesis. Integrative and Comparative Biology, 58(3), 567–577. 10.1093/icb/icy073

Jastroch, M., Divakaruni, A. S., Mookerjee, S., Treberg, J. R., & Brand, M. D. (2010). Mitochondrial proton and electron leaks. Essays in Biochemistry, 47, 53–67. 10.1042/bse0470053

Kikusato, M., & Toyomizu, M. (2013). Crucial Role of Membrane Potential in Heat Stress-Induced Overproduction of Reactive Oxygen Species in Avian Skeletal Muscle Mitochondria. PLOS ONE, 8(5), e64412. 10.1371/journal.pone.0064412

Koch, R. E., Buchanan, K. L., Casagrande, S., Crino, O., Dowling, D. K., Hill, G. E., Hood, W. R., McKenzie, M.,Mariette, M. M., Noble, D. W. A., Pavlova, A., Seebacher, F., Sunnucks, P., Udino, E., White, C. R., Salin, K., & Stier, A. (2021). Integrating Mitochondrial Aerobic Metabolism into Ecology and Evolution. Trends in Ecology & Evolution, 36(4), 321–332. 10.1016/j.tree.2020.12.006

Korshunov, S. S., Skulachev, V. P., & Starkov, A. A. (1997). High protonic potential actuates a mechanism of production of reactive oxygen species in mitochondria. FEBS Letters, 416(1), 15–18. 10.1016/S0014-5793(97)01159-9

Kuznetsova, A., Brockhoff, P. B., & Christensen, R. H. B. (2017). lmerTest Package:Tests in Linear Mixed Effects Models. Journal of Statistical Software, 82, 1–26. 10.18637/jss.v082.i13

Lenth, R., Singmann, H., Love, J., Buerkner, P. and Herve, M., 2019. Package ‘emmeans’. R package version, 1(3.2).

López-Lluch, G., Hunt, N., Jones, B., Zhu, M., Jamieson, H., Hilmer, S., Cascajo, M. V., Allard, J., Ingram, D. K., Navas, P., & de Cabo, R. (2006). Calorie restriction induces mitochondrial biogenesis and bioenergetic efficiency. Proceedings of the National Academy of Sciences of the United States of America, 103(6), 1768–1773. 10.1073/pnas.0510452103

Monaghan, P., Metcalfe, N. B., & Torres, R. (2009). Oxidative stress as a mediator of life history trade-offs:Mechanisms, measurements and interpretation. Ecology Letters, 12(1), 75–92. 10.1111/j.1461-0248.2008.01258.x

Mookerjee, S. A., Gerencser, A. A., Watson, M. A., & Brand, M. D. (2021). Controlled power:How biology manages succinate-driven energy release. Biochemical Society Transactions, 49(6), 2929–2939. 10.1042/BST20211032

Munro, D., & Treberg, J. R. (2017). A radical shift in perspective:Mitochondria as regulators of reactive oxygen species. Journal of Experimental Biology, 220(7), 1170–1180. 10.1242/jeb.132142

Neretti, N., Wang, P.-Y., Brodsky, A. S., Nyguyen, H. H., White, K. P., Rogina, B., & Helfand, S. L. (2009). Long-lived Indy induces reduced mitochondrial reactive oxygen species production and oxidative damage. Proceedings of the National Academy of Sciences of the United States of America, 106(7), 2277–2282. 10.1073/pnas.0812484106

Porter, R. K., & Brand, M. D. (1993). Body mass dependence of H+ leak in mitochondria and its relevance to metabolic rate. Nature, 362(6421), 628–630. 10.1038/362628a0

Quinlan, C. L., Treberg, J. R., Perevoshchikova, I. V., Orr, A. L., & Brand, M. D. (2012). Native rates of superoxide production from multiple sites in isolated mitochondria measured using endogenous reporters. Free Radical Biology and Medicine, 53(9), 1807–1817. 10.1016/j.freeradbiomed.2012.08.015

Remeŝ, V., & Martin, T. E. (2002). Environmental influences on the evolution of growth and developmental rates in passerines. Evolution, 56(12), 2505–2518. 10.1111/j.0014-3820.2002.tb00175.x

Roussel, D., Boël, M., & Romestaing, C. (2018). Fasting enhances mitochondrial efficiency in duckling skeletal muscle by acting on the substrate oxidation system. Journal of Experimental Biology, 221(4), jeb172213. 10.1242/jeb.172213

Roussel, D., & Voituron, Y. (2020). Mitochondrial Costs of Being Hot:Effects of Acute Thermal Change on Liver Bioenergetics in Toads (Bufo bufo). Frontiers in Physiology, 11. 10.3389/fphys.2020.00153

Salin, K., Auer, S. K., Rey, B., Selman, C., & Metcalfe, N. B. (2015). Variation in the link between oxygen consumption and ATP production, and its relevance for animal performance. Proceedings. Biological Sciences, 282(1812), 20151028. 10.1098/rspb.2015.1028

Salin, K., Luquet, E., Rey, B., Roussel, D., & Voituron, Y. (2012). Alteration of mitochondrial efficiency affects oxidative balance, development and growth in frog (Rana temporaria) tadpoles. Journal of Experimental Biology, 215(5), 863–869. 10.1242/jeb.062745

Salin, K., Roussel, D., Rey, B., & Voituron, Y. (2012). David and Goliath:A Mitochondrial Coupling Problem? Journal of Experimental Zoology Part A: Ecological Genetics and Physiology, 317(5), 283–293. 10.1002/jez.1722

Salin, K., Villasevil, E. M., Anderson, G. J., Lamarre, S. G., Melanson, C. A., McCarthy, I., Selman, C., & Metcalfe, N. B. (2019). Differences in mitochondrial efficiency explain individual variation in growth performance. Proceedings of the Royal Society B: Biological Sciences, 286(1909), 20191466. 10.1098/rspb.2019.1466

Salin, K., Villasevil, E. M., Anderson, G. J., Selman, C., Chinopoulos, C., & Metcalfe, N. B. (2018). The RCR and ATP/O Indices Can Give Contradictory Messages about Mitochondrial Efficiency. Integrative and Comparative Biology, 58(3), 486–494. 10.1093/icb/icy085

Schielzeth, H., Dingemanse, N. J., Nakagawa, S., Westneat, D. F., Allegue, H., Teplitsky, C., Réale, D., Dochtermann, N. A., Garamszegi, L. Z., & Araya-Ajoy, Y. G. (2020). Robustness of linear mixed-effects models to violations of distributional assumptions. Methods in Ecology and Evolution, 11(9), 1141–1152. 10.1111/2041-210X.13434

Sirsat, S. K. G., & Dzialowski, E. M. (2018). Ontogeny of skeletal and cardiac muscle mitochondria oxygen fluxes in two breeds of chicken. Comparative Biochemistry and Physiology Part A: Molecular & Integrative Physiology, 215, 20–27. 10.1016/j.cbpa.2017.10.017

Smith, S. M., Nager, R. G., & Costantini, D. (2016). Meta-analysis indicates that oxidative stress is both a constraint on and a cost of growth. Ecology and Evolution, 6(9), 2833–2842. 10.1002/ece3.2080

Speakman, J. R., Blount, J. D., Bronikowski, A. M., Buffenstein, R., Isaksson, C., Kirkwood, T. B. L., Monaghan, P., Ozanne, S. E., Beaulieu, M., Briga, M., Carr, S. K., Christensen, L. L., Cochemé, H. M., Cram, D. L., Dantzer, B., Harper, J. M., Jurk, D., King, A., Noguera, J. C.,…Selman, C. (2015). Oxidative stress and life histories:Unresolved issues and current needs. Ecology and Evolution, 5(24), 5745–5757. 10.1002/ece3.1790

Stearns, S. C., & Koella, J. C. (1986). The evolution of phenotypic plasticity in life-history traits:Predictions of reaction norms for age and size at maturity. Evolution, 40(5), 893–913. 10.1111/j.1558-5646.1986.tb00560.x

Stier, A., Bize, P., Hsu, B.-Y., & Ruuskanen, S. (2019). Plastic but repeatable:Rapid adjustments of mitochondrial function and density during reproduction in a wild bird species. Biology Letters, 15(11), 20190536. 10.1098/rsbl.2019.0536

Stier, A., Metcalfe, N. B., & Monaghan, P. (2020). Pace and stability of embryonic development affect telomere dynamics:An experimental study in a precocial bird model. Proceedings of the Royal Society B: Biological Sciences, 287(1933), 20201378. 10.1098/rspb.2020.1378

Stier, A., Monaghan, P., & Metcalfe, N. B. (2022). Experimental demonstration of prenatal programming of mitochondrial aerobic metabolism lasting until adulthood. Proceedings. Biological Sciences, 289(1970), 20212679. 10.1098/rspb.2021.2679

Vaillancourt, E., Prud’Homme, S., Haman, F., Guglielmo, C. G., & Weber, J.-M. (2005). Energetics of a long-distance migrant shorebird (Philomachus pugnax) during cold exposure and running. Journal of Experimental Biology, 208(2), 317–325. 10.1242/jeb.01397

